# Looming stimuli reliably drive innate, but not learned, defensive responses in rats

**DOI:** 10.1101/2022.02.07.479432

**Authors:** Mirjam Heinemans, Marta A. Moita

## Abstract

Survival relies on an organism’s intrinsic ability to instinctively react to stimuli such as food, water, and threats, ensuring the fundamental ability to feed, drink, and avoid danger even in the absence of prior experience. These natural, unconditioned stimuli can also facilitate associative learning, where pairing them consistently with neutral cues will elicit responses to those cue. Threat conditioning, a well-explored form of associative learning, commonly employs painful electric shocks, mimicking injury, as unconditioned stimuli. It remains elusive whether actual injury or pain is necessary for effective learning, or whether the threat of harm is sufficient. Approaching predators create looming shadows and sounds, triggering strong innate defensive responses like escape and freezing. This study investigates whether visual looming stimuli can induce learned freezing or learned escape responses to a conditioned stimulus in rats. Surprisingly, pairing a neutral tone with a looming stimulus only weakly evokes learned defensive responses, in contrast to the strong responses observed when the looming stimulus is replaced by a shock. This dissociation sheds light on the boundaries for learned defensive responses thereby impacting our comprehension of learning processes and defensive strategies.

## Introduction

Most of our knowledge regarding survival circuits in the brain comes from threat conditioning studies^1–3^. These studies mainly focus on defensive behavioral responses, such as freezing, to learned cues predicting an aversive stimulus. Electric shocks, which may mimic painful injury upon contact with a predator, are predominantly used as the unconditioned aversive stimulus. More recently, other noxious stimuli have been shown to effectively drive this form of learning, such as heat^4–6^, or activation of pain responsive brain regions^7^. Non-painful threatening stimuli are seldomly used for threat learning, even though animals can rapidly detect danger through visual, chemical and auditory cues, and avoid contact through rapid innate defensive responses like freezing and escape^8^. Whether these cues (artificial cues similar to those produced by the presence or approach of a predator) are able to drive learning remains unclear. Predator odors have been used to drive threat learning with variable results. In most cases the context in which subjects were exposed to predator odors fails to drive freezing or escape responses, although sometimes it drives avoidance behaviors^9–11^. One possibility is that a predator’s odor, often present in excretions, does not imply the predator’s proximity, thus constituting a remote cue of threat. Hence, animals may learn to avoid the location where the odor was scented, but will not learn to exhibit acute defensive responses like escape or freezing, typically triggered by an imminent encounter with a predator.

A looming stimulus, in the form of a rapidly expanding black disk that mimics the shadow of an approaching predator, elicits strong innate freezing and escape responses in virtually all visual animals tested in the lab, to varying degrees, including invertebrates and vertebrates such fish, rodents and humans^12–18^. The immediacy of threat associated with a looming stimulus may result in animals learning more effectively to associate cues and locations with its presence and to display acute defensive responses to these cues. Studies in mice have shown that the superior colliculus, where information about visual looming stimuli is processed, sends projections to the dorsal periaqueductal gray (dPAG) capable of driving escape responses. The superior colliculus also projects to the amygdala, which is required for looming-triggered freezing^19–21^. This looming-triggered freezing is possibly driven by projections from the amygdala to the ventral PAG – a brain structure essential for threat learning^1,3,22^.

To test the efficacy of looming stimuli in driving threat learning, we developed a conditioning protocol where a neutral pure tone, the conditioned stimulus, was either paired with foot shock or with a visual looming stimulus that robustly induces freezing and escapes. It was previously demonstrated that looming stimuli do not efficiently drive contextual threat learning^18^, therefore, in this study, we chose a cued conditioning task that typically drives conditioned responses more reliably than contextual conditioning paradigms. Specifically, we used a tone-shock or a tone-loom paradigm in which a neutral tone cued the arrival of electric shock or looming shadow. We used these conditioning protocols in two separate experiments – one focused on freezing, the other on escape – to assesses whether rats exhibit either of these acute defensive behaviors in response to tones associated with visual looms.

## Results

### Looming stimuli are weak drivers of learned freezing responses

The first experiment was performed to assess the amount of learned freezing triggered by a tone previously associated to either a shock or a visual loom. Rats were put in a conditioning box and exposed to a neutral tone followed by either a foot shock or a visual looming stimulus (fig. 1A). In the conditioning session of this experiment, rats in both groups displayed similarly low freezing during the baseline period (median freezing of shock condition: 1.87%; median freezing of loom condition: 4.35%; U=251, p=0.92) and showed equivalently high levels of freezing during the tone-shock or tone-loom pairings (fig. 1B, median percentage increase in freezing upon exposure to tone-shock pairings: 64.76% and upon tone-loom pairings: 53.63%; U=56, p=0.65; see supplemental figure 1 for a similar analysis focusing on freezing during tone presentations). The following day both groups of rats were tested for their ability to display learned freezing to the tone cue in a neutral environment. Again, little or no freezing was observed during the baseline period (shock: 0.22% freezing, loom: 0.51% freezing). However, upon the presentation of the tone cue, rats previously exposed to tone-shock pairings during conditioning showed robust freezing that was significantly higher than the low freezing displayed by rats previously exposed to tone-loom pairings (fig. 1C and supplemental videos 1 (shock) and 2 (loom), median increase in shock condition: 94.27%; median increase in loom condition: 5.86%, U=2, p<0.0001). Figure 1B shows a small decrease in freezing at the end of the training session, in response to the last few tone-loom presentations, a decrease that seems absent in the tone-shock group. This could reflect the beginning of extinction and explain the lack of robust conditioned responses by rats in the tone-loom group to the tone in the test session. To investigate this possibility, we quantified freezing to each tone during the training session and verified that indeed there was a decrease in the tone-loom group. This decrease was not present in all animals in the tone-loom group (see supplemental figure 2A). Therefore, we next analyzed the behavior of rats that sustained high levels of freezing throughout the training session (suppl. fig. 2B and C). We found that these rats also showed weak responses to the tone in the test session (suppl. fig. 2C, median increase = 5.86% for tone-loom, W=0, p=0.0076), suggesting that extinction during training is unlikely to explain the low levels of conditioned freezing in rats exposed to tone-loom pairings.

**Figure 1.**
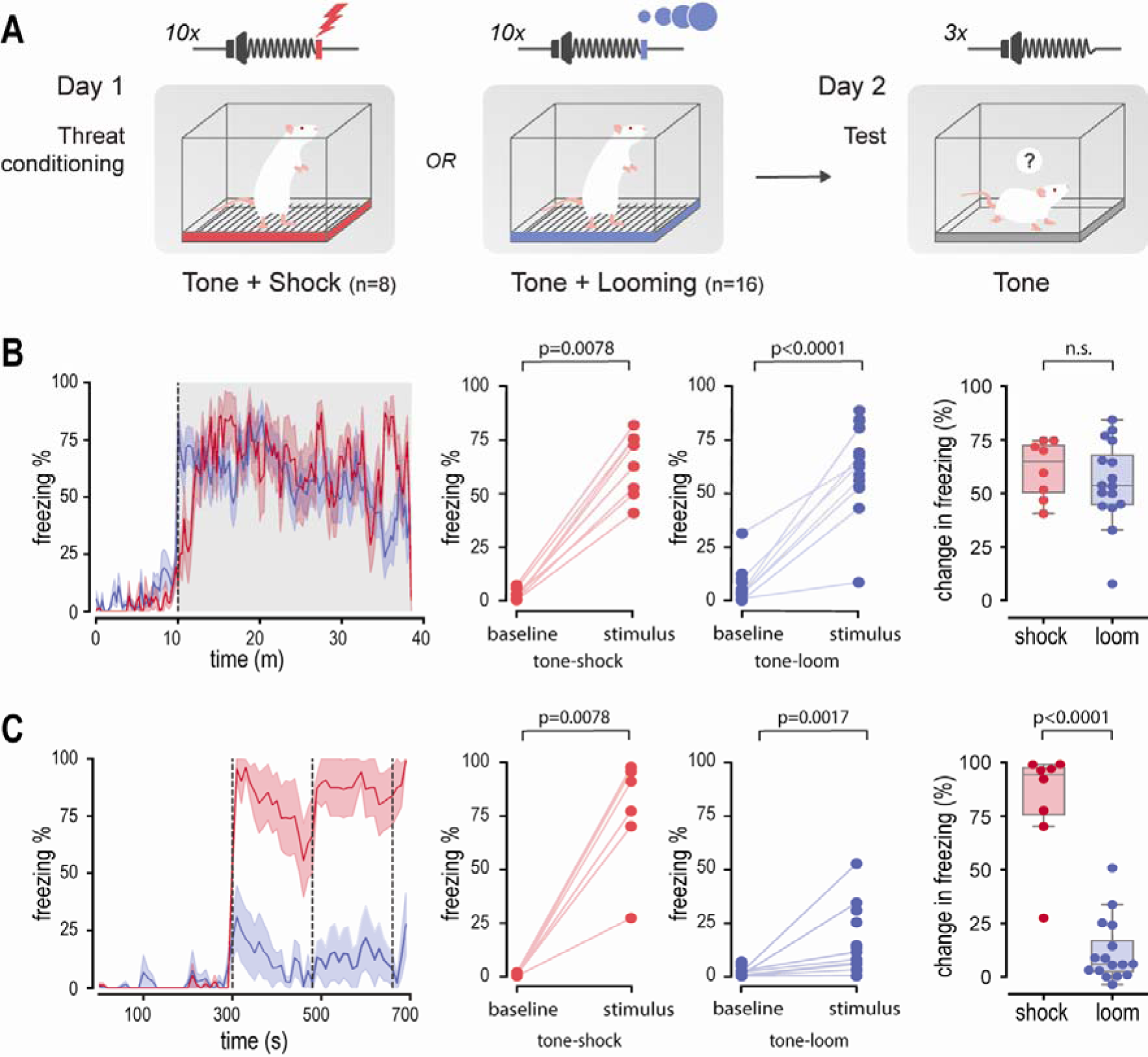
Rats freeze more to shock-associated tones than loom-associated tones. **(A)** Diagram of behavioral paradigm used to study learned freezing response to conditioned neutral tone. Rats received 10 tone-shock or tone-loom pairings and the next day exposed to 3 tone presentations. **(B)** Freezing during threat conditioning. Left: percentage of time freezing throughout the training session per ten-second epoch. Baseline period in white (0 until 10 minutes) and period of exposure to tone-shock/tone-loom pairings in gray (10 until 40 minutes). The line depicts average and the shade represents SEM. Middle two panels: the total percentage of freezing during baseline and stimulation presentation period per animal, for tone-shock and tone-loom conditions respectively. Right: change in percentage of time freezing (stimulus presentation period – baseline). Each dot corresponds to an individual animal; the box and whiskers represent the median and interquartile ranges. **(C)** Same as in **(B)** for freezing during the recall test. Gray dotted vertical lines indicate tone delivery. In all panels rats exposed to tone-shock pairings are depicted in red and rats exposed to tone-loom pairings in blue.

Although the freezing response to loom-associated tones was low, it was not absent, as the increase in freezing from baseline to the tone was statistically significant (fig. 1C, median increase in freezing: 5.74%, W=5, p=0.0017). The weak freezing responses to the tone after conditioning with the looms (median = 6.98% of time spent freezing), whether the result of a weak associative learning process or the response to the salient tone (well known to drive weak freezing responses, as those observed here), pales in the face of the robust responses to the tone after conditioning with shocks (median = 94.5%). If the weak response to the tone in the test session is driven by stimulus saliency, it should not be different from the first tone presentation of the training session, before rats were exposed to either a shock or a loom. Indeed, we found no difference in freezing responses to the first tone presentation of the training and test sessions for rats in the tone-loom group, and a significant increase in this response for rats in the tone-shock group (suppl. fig. 1F; for tone-loom median increase is 4.83%, W=21, p=0.16; for tone-shock the median increase is 33.4%, W=4, p=0.027). Furthermore, tone-evoked freezing responses decreased over the course of the test session for rats in the tone-loom group, while rats in the tone-shock group increased this response (fig.1C and suppl. fig. 1D). In sum, these results indicate that a visual looming stimulus, while a potent driver of innate freezing, only weakly drives freezing in response to a previously neutral cue associated with it, a response possibly caused by the stimulus saliency.

### Looming stimuli are weak drivers of learned escape responses

To test conditioned escape responses, we adapted an existing task where rats display escapes in response to an approaching naturalistic predator^23^. We modified this task to test escapes in response to a visual looming stimulus or to a conditioned tone that was previously paired either with shocks or looms. Rats were first trained during two consecutive days to retrieve food pellets placed at increasing distances; up to 1 meter from a shelter located at one end of a 2-meter runway (fig. 2A). Most rats slowly came out of the shelter and ran back to it as soon as they got the pellet, consuming the pellet inside the shelter (fig. 2B), suggesting that indeed they regarded the shelter as their “safe space” in the setup.

**Figure 2.**
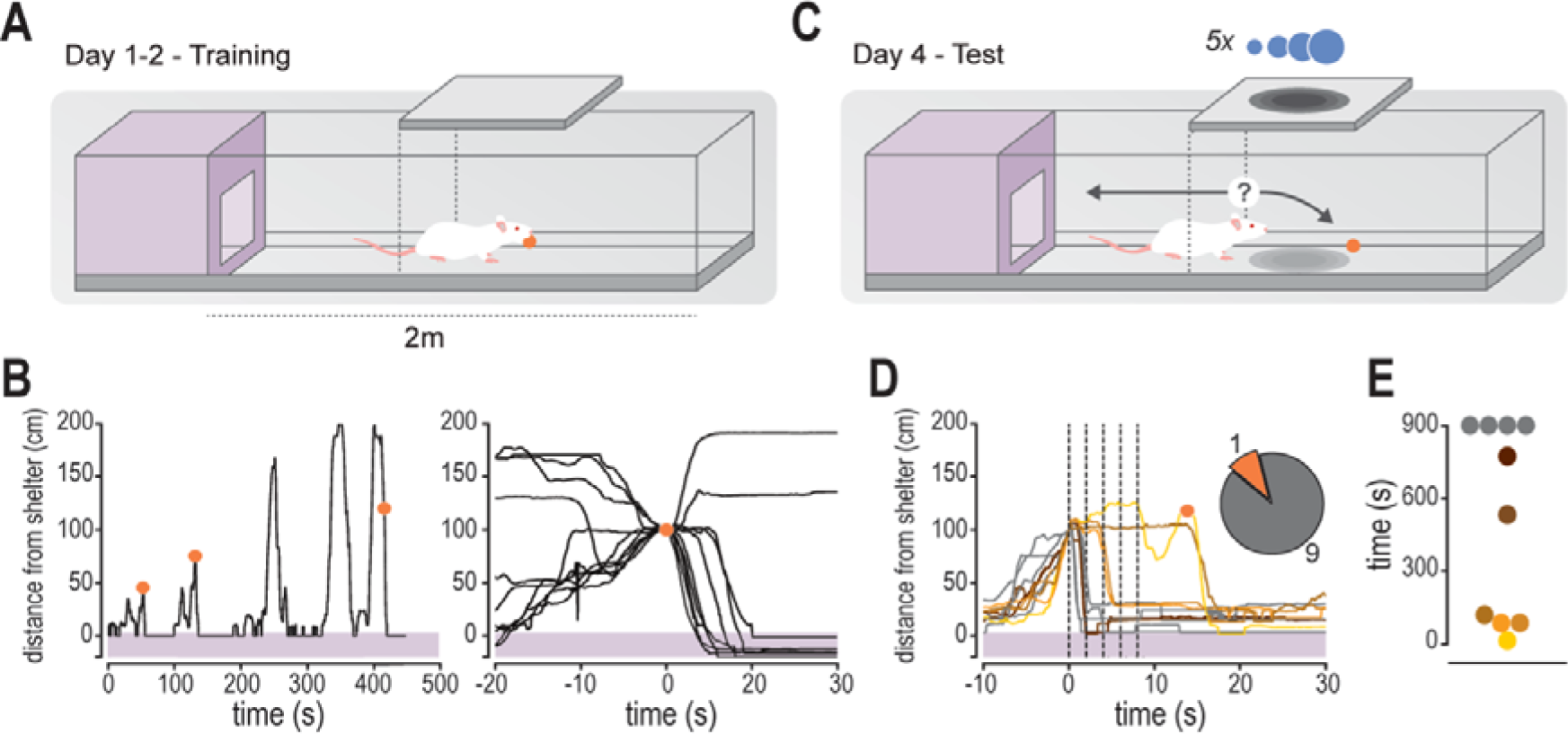
Escape responses upon looming stimulus presentation during pellet retrieval task. **(A)** Diagram of runway used in pellet retrieval training sessions. Rats were trained to retrieve yoghurt pellets on two consecutive days. Pellets were placed at an increasing distance from the shelter on the left: 0.25, 0.5 and 0.75m from shelter entrance on the first day, 0.5, 0.75 and 1.0m distance on the second day. **(B)** Left: example trace of a rat that retrieved 3 pellets within 500 seconds. The purple shaded area indicates shelter location (-30 to 0 cm) and orange dots indicate when and where a pellet was retrieved. Right: the traces of all animals (n=10) aligned to retrieval of the last pellet (time of pellet retrieval = 0s). All but two rats went back to the shelter once they retrieved the pellet. **(C)** Diagram of test day. 5 consecutive looms are triggered when rats reached a virtual threshold distance from the shelter (90cm, represented by the dashed line). **(D)** Trajectory of rats aligned to time of loom presentations indicated by vertical dashed lines, starting at t=0s. Most rats returned to the shelter after the first loom, two after the second loom, and only one rat retrieved the pellet before returning to the shelter (orange dot). Inset, pie chart indicating number of pellets retrieved before re-entering the shelter. **(E)** Time elapsed between crossing of loom triggering threshold and pellet retrieval. Each dot corresponds to one animal. Lighter dots represent shorter times for pellet retrieval, while darker colors represent longer retrieval times. Gray dots represent rats that failed to retrieve the pellet within 15 minutes after loom presentation. The colors of the dots are matched to that of the trajectories of the same animals in **(D)**.

After the two days of pellet retrieval training, we tested whether looms provoked escape to the shelter in this setup. Rats were tested in the runway with a pellet 1 meter away from the shelter. Once rats reached the middle of the runway, 10 cm before the pellet’s location, a train of looming stimuli was presented on an overhead screen (totaling five 0,5s looms with a 1s interval, fig. 2C). Of the 10 rats tested, one retrieved the pellet before running back to the shelter, with all other 9 rats escaping to the shelter without the pellet (fig. 2D, supplemental video 3). Of these, 5 rats exited the shelter to retrieve the pellet in the remaining 15 minutes of the experiment (requiring varying times to do so), while the other 4 rats failed to retrieve the pellet (fig. 2E). Interestingly, the rats that took a shorter time to retrieve the pellet were also the ones that required repeated presentation of the looming stimuli to flee to the shelter (fig. 2D and 2E). Hence, both an empty-handed return to the shelter and the amount of time taken to retrieve the pellet seem to be good indicators of the perceived threat level.

Having established the loom-triggered escape task, we turned to studying learned escape responses. We conditioned a new set of rats to a tone cue paired with either shocks or looms, as in the experiment before (except the tone was 1 second long, see Methods and fig. 2A), and tested their response to the tone in the pellet retrieval task. Whereas the lack or low level of freezing responses to a neutral tone has been extensively reported, the response of rats to a neutral yet salient stimulus in this pellet retrieval task was not known, thus, we added a tone-alone control group (see Methods). All of the tone-shock conditioned rats escaped to the shelter before retrieving the pellet (8/8 rats), whereas in the tone-loom and the tone alone groups only 3/7 and 3/6 rats, respectively, did so (fig. 3B, supplemental videos 4-6 for tone-shock, tone-loom, and tone conditions respectively). In addition, rats in the tone-shock group took longer to retrieve the pellet than rats in both other groups, which showed similar pellet retrieval times (fig. 3C; we used “pellet survival” curves, which take into account whether and when a pellet was retrieved during the test session; Chi square=11.00, p-value = 0.004). It is possible that the difference seen in escape responses between the tone-shock and tone-loom groups stems from the lower conditioning levels to the tone of the latter group, reflected in lower freezing levels observed during conditioning of these rats (see supplemental figure 3). However, none of rats in the tone-shock group retrieved the pellet before retreating to the shelter, regardless of their freezing levels (which varied between 42 and 87%, suppl. fig. 3A). In addition, we observed no clear relationship between pellet retrieval time and amount of freezing during conditioning in any of the groups (suppl. fig. 3C).

**Figure 3.**
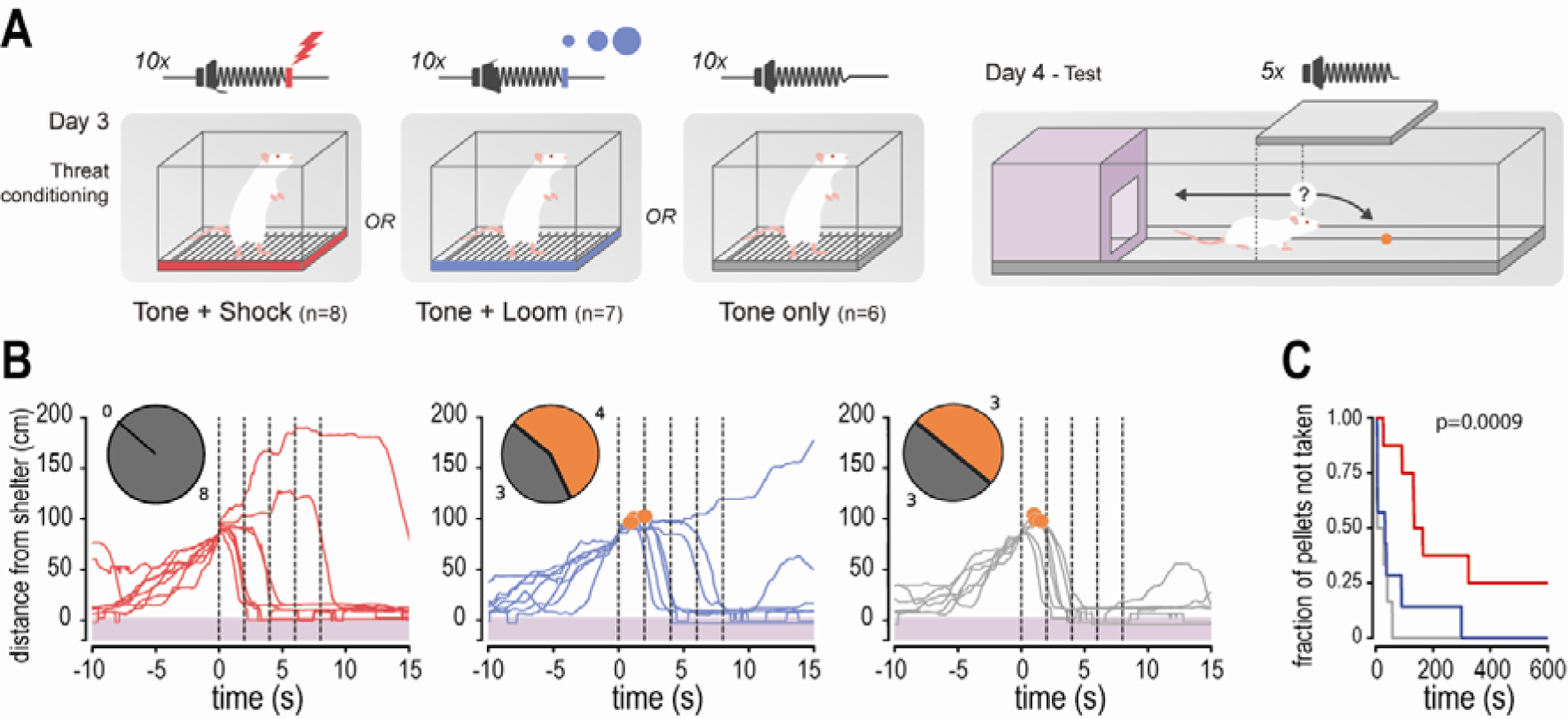
Escape responses upon conditioned tone presentation during and pellet retrieval task. **(A)** Behavioral paradigm to test conditioned escapes after tone-shock or tone-loom exposure. After two days of training the pellet retrieval task, rats received ten presentations of a neutral tone paired with a shock, a visual loom, or nothing. The next day (day 4), the rats were placed in the runway with a pellet located at a 1m distance from the shelter. Crossing the virtual threshold in between the shelter and pellet (dashed line) triggered the presentation of 5 consecutive pure tones previously associated to shock or loom, or neutral. **(B)** Trajectory of rats aligned to time of tone presentations indicated by vertical dashed lines, starting at t=0s. Orange dots indicate when and where a pellet was retrieved. Inset, pie chart indicates number of pellets retrieved before re-entering the shelter. **(C)** “Pellet survival” plots in the different conditions. The p-value represents significance of the differences in survival of the pellets as calculated by a Kaplan-Meier test. The significant difference is driven by the increased survival rates of the pellets in the tone-shock group compared to the two remaining groups. In all panels, rats exposed to tone-shock pairings are depicted in red, rats exposed to tone-loom pairings in blue and rats exposed to tone alone in gray.

The remaining behaviors, including rearing, reaching, scanning and freezing were scored, but there were no differences in those behaviors across conditions (data not shown). Taken together, these results show that, similar to learned freezing responses, looming stimuli are not effective at driving robust learned escapes, despite evoking robust innate escape responses.

## Discussion

Here we establish that a visual looming stimulus, a feature of an approaching predator, induces robust innate defense responses in rats. Using two different setups, we show that if a shelter is available, male rats escape to the shelter upon seeing visual looms, while they freeze to looming stimuli if no shelter is present. This indicates that the rats’ choice between freezing and fleeing is heavily modulated by context, as previously reported in mice^16,24^. Yet, an auditory cue repeatedly paired with the same looming stimulus failed to trigger these behaviors in a robust manner, unlike auditory cues paired with shock that drive both responses strongly (fig. 2C and 3C).

Further experiments are required to establish the boundaries of looming triggered learning, including investigating the behavior of female rats, which show defensive responses distinct to those of males in response to the same threatening stimuli^25–27^. Changing the features of the looming stimuli, for example using faster looms, changing the properties of the conditioned cues, either using different sounds, for example natural sounds, or even other sensory modalities, and/or looking at other behaviors such as avoidance, might provide useful insights. Still, the discrepancy between the ability of looming stimuli to drive innate responses and the ability to drive these same responses through association with other cues is evidenced in this study. In addition, it was previously shown that exposure to multiple looming stimuli failed to drive contextual threat learning^18^. Crucially the two studies differ in both the looming stimuli used (here, each tone cue was paired with a single loom, with an average ITI of 3 minutes, in the previous study 8 trains of 20 looms delivered at 1Hz, with an ITI of 30s were used) and in the conditioned stimulus (here a pure tone cue was used, in the previous study, contextual cues, the conditioning box, were used).

Multiple factors may explain the discrepancy observed. Looming shadows and foot shocks may activate neuronal circuits that are distinct from those that drive threat learning and control the expression of freezing and escapes in responses to learned cues. Although it has been proposed that innate and learned defensive responses rely on partially distinct circuits^1^, more recent studies suggest considerable overlap of involved brain areas. The lateral (LA), basal (BA) and central (CeA) nuclei of the amygdala, are crucial for threat learning and conditioned freezing^3,28,29^. Of these, the LA and CeA have also been implicated in looming triggered freezing^30^ or escape followed by freezing^31,32^. Strikingly, auditory cues paired with optogenetic activation of the Superior Colliculus (SC), involved in the processing of visual looms lead to conditioned freezing, indicating that a sufficiently strong activation of the SC could evoke learned freezing responses^31^. In addition to conditioned freezing, rats also show conditioned escapes in response to cues predicting shocks if given the opportunity. Active avoidance is thought to be predominantly processed in the LA and BA, and then relayed to the Nucleus Accumbens^33–36^. Similar to conditioned escapes, inactivating or lesioning the amygdala of rats abolished innate escape and avoidance responses to an artificial robotic predator in a naturalistic foraging task^23^. More recently, it has been shown that different sub-populations of cells in central amygdala drive freezing and avoidance responses upon socially triggered threat^37^. However, Evans et al.^38^ showed that innate escapes to looming stimuli can bypass the amygdala, through direct projections from SC to the dorsal periaqueductal gray, where escape responses are thought to be initiated^39^.

Finally, another circuit involving dopaminergic projections to the tail of the striatum has been implicated in both innate and learned avoidance^40^. In summary, looming stimuli can activate various circuits, most of which include sub-nuclei of amygdala widely implicated in the expression of learned freezing and escape responses.

It is also possible that only close encounters with predators, resulting in injury or pain, lead to a learning process capable of driving acute defensive responses, such as freezing or escape, in response to cues associated with the encounter. Information about pain enters the brain through parallel ascending pathways reaching various sub-nuclei of the amygdala the LA, BA and CeA^7,41^. Silencing either the parabrachial nucleus (PBN) or the dorsal periacqueductal gray (dPAG), which provide pain information to the amygdala, during conditioning attenuates conditioned freezing responses^7,41^, while pairing a neutral tone with activation of the PBN or dPAG is sufficient to drive learned freezing to the tone cue in rodents^7,42^. These findings illustrate the importance of pain information reaching the amygdala during threat learning. Experiencing a painful stimulus evokes the release of neuromodulators such as noradrenaline and acetylcholine, which modulate activity in the amygdala^43–47^. Decreasing or increasing noradrenergic or cholinergic modulation of the amygdala has been shown to attenuate or enhance threat learning, respectively^48–50^. Given that the SC projects directly to the dPAG and the dPAG in turn projects to the locus coeruleus (LC), the main source of noradrenaline to the amygdala, looming stimuli could in theory trigger a noradrenergic response similar to the one triggered by shocks^38,51,52^. However, it is currently unknown whether the SC has a functional connection to the LC through the PAG. To examine whether looming stimuli could induce the release of neuromodulators to a similar extent as shocks, more research is required.

Defensive responses may vary depending on proximity to threat^53^. Upon inevitable contact with a threat, rats show a burst of activity or fight. Upon threat detection, either more distant or when escape is possible, rats will show acute defensive responses, respectively freezing and escape. If threat is more remote, such as when the presence of a predator is likely but not detected yet, other responses such as avoidance or risk assessment may be displayed. Indeed, shocks, mimicking contact with the threat, typically drive the unconditioned responses jumping, running, and squeaking, whereas cues associated with shock, thus one step removed from the threat, typically drive freezing or escape if possible^54,55^. A looming stimulus, an innate cue of impending threat, can also be considered one step removed from the threat, and just as learned cues of shock it drives freezing and escape. It follows that cues associated with the loom represent an even more distant threat, which could be learned through second order conditioning. This could lead to a different set of defensive behaviors such as risk assessment and avoidance, or weaker learned freezing or escapes. Although we did not find evidence of learned freezing or escape, beyond the weak responses observed to the novel salient tone, future studies are needed where one would vary a number of parameters, such as looming speed, number of pairings, inter-trial intervals, size of arenas, etc.

The fact that we did not see robust learned defensive responses does not necessarily mean that the rats did not learn about the relationship between tone and loom. For example, sensory pre-conditioning studies show that rats can learn to associate two neutral stimuli with each other, but this learning is not accompanied by any overt behavioral response^56,57^. The same might be true in our study: rats may learn about the association between the tone and the looming stimulus, but not show an acute, overt, defensive response to the tone. Visual looming stimuli, often generated by an approaching predator, may fail to evoke acute defensive responses after being paired with a tone. However, the detection of a predator, involving multisensory integration, could be more effective at driving learning resulting in freezing or escapes triggered by neutral cues associated to the predator, even in the absence of a painful interaction.

In conclusion, our work broadens our understanding of learned defensive responses and their boundaries, by showing that the detection of innate cues of threat leads to robust acute defensive responses, but these cues by themselves are not as effective drivers of threat learning. Several requirements may have to be met for a stimulus to be able to drive learned acute defensive responses. An acute defensive response, such as escape or freezing, may require proximity of the threat, while other responses such as avoidance or risk assessment may be displayed in response to more distant threats^53^. A learned cue, i.e., a stimulus that is associated with the presence of the threat, is necessarily at least one step further from the threat itself. Indeed, foot shocks typically drive the unconditioned responses jumping, running, and squeaking, whereas cues associated with shock typically drive learned freezing ^55^. Arguably freezing is a response adjusted to a more distant threat than jumping and squeaking. It follows that while a looming stimulus drives freezing and escape, cues associated with the loom represent an even more distant threat, possibly driving avoidance. This implies that for an acute defensive response to be displayed upon the presentation of a learned cue, the learned association between the cue and the threat must be very strong which in turn depends on the degree of the threat, such as those that result in pain or injury. However, whether these constraints of threat learning hold for wild rats inn their natural habitat, needs to be tested. Still, the findings in this study suggest that the emotional state of an animal in danger can be quite distinct for different kinds of threat, even when the overt behavioral response is similar, opening the path for the mapping between features of emotional states of fear and the forms of learning they afford. The dissociation between the capacity to drive an innate defensive response and to drive threat learning raises new questions regarding the functioning of survival mechanisms.

## Materials & Methods

### Subjects

A total of 60 naïve male Sprague Dawley rats, weighing 225g to 250g, were obtained from Charles Rivers Laboratories (France). All animals used were males as all our prior experiments, protocols etc., were optimized for male rats and there are sexual dimorphisms ^25–27^. Upon arrival, the animals were pair-housed in Plexiglas top filtered ventilated cages (GR900 for rats, Tecniplast S.p.A, Italy) with *ad libitum* access to water and food. They were maintained on a 12h light/dark cycle (lights off at 8 p.m.), a temperature of 20-22°C and 40-70% humidity. After a one-week acclimatization, the experimenter handled all animals on three consecutive days in the week preceding experimental procedures. All animal procedures were performed under the guidelines of the Animal Welfare Body of the Champalimaud Research (Portugal) and in strict accordance with the European Community’s Council Directive (86/609/EEC).

### Behavioral apparatus

#### Shock-conditioning

The shock conditioning box (model H10-11RTC, Coulbourn Instruments; 30.5cm width x 30.5cm height x 25.4cm depth) was equipped with a metal grid floor to deliver foot shocks (model H10-11RTC-SF, Coulbourn Instruments) and placed inside a sound isolation chamber (Action, automation and controls, Inc) with white walls. The side walls of the conditioning box were made of clear Plexiglas and cleaned with rose scented detergent after every conditioning round. A precision programmable shocker (model H13-16, Coulbourn Instruments) was used to deliver foot shocks. Pure tones (5kHz, 60dB) were produced by a sound generator (RM1, Tucker-Davis Technologies) and delivered through a horn tweeter (model TL16H80HM, VISATON). The rats’ behavior was tracked by a video camera mounted on the ceiling of each sound attenuating chamber. An infrared surveillance video acquisition system was used to record and store all videos on a hard disk and freezing behavior was scored manually offline.

#### Loom-conditioning

The visual loom conditioning box was made of a black acrylic floor with clear, dark red sides (30cm width x 50cm height x 55cm depth) and was cleaned with a lemon-scented detergent solution. This box was placed in a room with ceiling lights on. Pure tones were delivered through the same system as the shock conditioning (see above). Visual stimuli were projected with an LED projector (ML750e, Optoma Europe ltd, United Kingdom) onto an opaque white Plexiglass screen placed on top of the behavioral box. The behavior was captured with an infrared camera (PointGrey Integrated Imaging Solutions GmbH, Germany) and stored on a hard disk for offline manual scoring. Both the tone-loom delivery and the video capture were controlled by a custom workflow using the Bonsai visual programming language (Lopes et al., 2015).

#### Recall-test for freezing

The box to test conditioned freezing consisted of a chamber made of clear Plexiglas walls (30cm width x 34cm height x 27cm depth, Gravoplot). The floor contained a removable tray with bedding (the same used in the animals’ home cages). The box and tray were cleaned with water and 70% ethanol. The box was placed inside a sound attenuation chamber (80cm width x 52.5cm height x 56.5cm depth) made of MDF lined with high-density sound attenuation foam (MGO Borrachas Tecnicas) and a layer of rubber. Pure tones were delivered through the same equipment as described for conditioning (see above). The behavior of the animals was tracked by infrared video cameras mounted on the walls of the sound attenuating chambers. A surveillance video acquisition system was used to record and store all videos, and freezing behavior was scored using the FreezeScan software from Clever Sys. In all tests, the rats were considered to be freezing if they did not show any movement except breathing for at least one second.

#### Escape runway

A large runway (200cm length x 50cm width x 50cm height), with an adjacent shelter (30cm length x 50cm width x 50cm height) was used to look at escape behavior. The shelter consisted of a black acrylic floor, with red-transparent acrylic walls. A black acrylic plate was used as a roof, and could be removed to place rats in the shelter at the beginning of each experiment. The shelter was connected to the large runway through a small gate (10cm x 10cm) that could be closed with a transparent acrylic sliding door. The runway’s back and side walls, as well as the floor, were made of (waterproof) black painted wood. The front wall was made out of red-transparent acrylic, allowing for video recording of the rats’ behavior inside of the runway, and the ceiling consisted of transparent acrylic with white baking paper on top that functioned as a screen for the looming stimuli. Looming stimuli were projected with an LED projector (ML750e, Optoma Europe ltd, United Kingdom), and pure tones were delivered with a horn tweeter (model TL16H80HM, VISATON). The runway had two infrared lights at each far-end side illuminating the area, while the shelter had one on top. The behavior was captured by infrared cameras (PointGrey Integrated Imaging Solutions GmbH, Germany) and stored for later use. There was one sideways camera capturing the behavior of the rats in the large runway, and one filming the behavior in the shelter from above.

## Behavioral procedures

### Conditioned freezing experiment

#### Habituation and tone conditioning

On days 1-3 all rats were randomly exposed to one environment per day: the test box, the loom box, and the shock box, for 20 minutes. Afterwards, the animals were randomly assigned to either the tone-loom or tone-shock conditioning group. On day 4, rats in the tone-shock group were placed in the shock conditioning box, where they received 10 tone-shock pairings after a ten-minute baseline. The pure tones (5kHz, 60dB) lasted 10 seconds, immediately followed by a shock of 0.5mA lasting 0.5s. The interval between tone-shock presentations ranged from 1 to 5 minutes, with an average of 3 minutes. The animals in the tone-loom conditioning group were placed in the loom box, where they received 10 tone-loom pairings after a ten-minute baseline period. The tone had the same properties as that of the tone-shock pairings and was immediately followed by the looming stimulus: a black disk that increased exponentially from 1cm to 30cm in 1 second (l/v = 16.7ms). Again, inter-trial intervals varied between 1 and 5 minutes (average of 3 minutes). The behavior of all rats was recorded for offline scoring of freezing, which was calculated by taking the percentage of freezing per ten-second epoch throughout the duration of the conditioning.

#### Conditioned freezing test

The day after conditioning, all animals were placed in the test chamber individually, and after a 5-minute baseline they were exposed to three tones (same as described above) with a 3-minute inter-trial interval. The behavior of the rats was recorded for offline scoring of freezing using FreezeScan. Similar to the conditioning, freezing was calculated by taking the freezing percentage per ten-second epoch.

### Escape experiment

#### Habituation

Each day after handling (3 minutes for 3 days in the week prior to the start of the experiment), rats received yoghurt-flavored pellets (mini yoghurt drops, BioServ, United States) in their home cage, to habituate them to the treat. Prior to the beginning of the experiment, rats were habituated to the shelter. Habituation was achieved by placing the rats in the shelter with 3 yoghurt pellets for two consecutive days, and allowing twenty minutes for rats to explore the closed shelter and eat the pellets. On the first day of the experiment (training day 1) a single yoghurt pellet was placed in the runway, 25cm away from the shelter exit. After putting the rat in the shelter, the door to the runway was opened, and the rat was free to explore the runway and retrieve the pellet. As soon as the first pellet was retrieved and consumed, the sliding door was closed again while the rat was in the shelter, and a second pellet was placed at a 50cm distance from the shelter. The rat was once again allowed to explore and retrieve the pellet, and the sequence was repeated to place a third pellet at a distance of 75cm. The training session ended as soon as the animal had retrieved all three pellets, or if the total time of the session had reached 30 minutes. The second training session, the following day, was identical to the first, with the exception that the pellets were placed at a distance of 50, 75, and 100cm from the exit of the shelter. All training sessions were recorded with the Bonsai visual programming language (Lopes et al., 2015) and saved for offline analysis.

#### Conditioning

On day 3 of the experiment, the rats were assigned to either a loom-alone, tone-shock, tone-loom, or tone-alone condition. Loom-alone rats, used to test innate escape responses the looming stimuli, were put in the shock conditioning box for the same length of time as all other animals, but no stimuli were presented. The conditioning protocol for the tone-shock and tone-loom groups was performed as described for the conditioned freezing experiment (tones were 5kHz at 60dB; shocks were 0.5mA lasting 0.5s; looms increased exponentially from 1cm to 30cm in 1 second). In short, after a 10-minute baseline, animals were either exposed to ten tone-shock pairings, or ten tone-loom pairings, with an interval between 1 and 5 minutes (3-minute average). The animals in the tone-alone condition were placed in the shock conditioning box, and received ten pure tone presentations. The interval between the tones was the same as described for the other groups. The behavior of the animals was recorded and saved for offline scoring of freezing. In this experiment, the tones were 1 second rather than 10 seconds long, as a pilot experiment showed that with a 10-second conditioned tone, animals had time to retrieve the pellet and reach the shelter before the end of the tone (when the unconditioned stimulus was expected), whereas the 1-second tone did not allow for this.

#### Escape test

The day after the conditioning - day 4 - a single yoghurt pellet was placed in the runway at 100cm from the shelter. Afterwards, rats were placed in the shelter and the door to the runway was opened, allowing rats to freely explore the arena, like the pellet retrieval training sessions on days 1 and 2. However, this time rats triggered a stimulus as soon as they reached a 90cm distance from the shelter. For the rats in the loom-alone group, this stimulus was a series of five visual looming stimuli, with an inter-stimulus interval of 1 second. The looming stimuli expanded from 1cm to 30cm in 1 second. The rats in the three remaining groups received a series of five 1 second pure-tone stimuli (5kHz), with a 1s inter-stimulus interval. The automatic triggering of the stimuli and the recording of the behavior was done using the Bonsai visual programming language (Lopes et al., 2015).

### Criteria for animal exclusion

For the freezing experiment, there were a total of 10 rats in the tone-shock condition, and 18 in the tone-loom condition. In each condition, 2 animals were excluded due to high freezing levels during the baseline, making it impossible to infer the effect of the conditioned tone on their freezing behavior. The behavior of 8 and 16 animals respectively was analyzed. For the escape experiment, 32 animals were used in total. Of those, 7 rats did not manage to retrieve all three pellets on training day 2, and 1 rat managed to open the sliding door by itself during the test, leading to the exclusion of these 8 animals.

### Video analysis

The freezing behavior during the cue test of the conditioned freezing experiment was scored automatically using FreezeScan from Clever Sys. This requires optimization and validation, which depends on the cameras used, size of the boxes and illumination, which was previously done for the cue testing chambers (Pereira, Cruz, Lima, & Moita, 2012). Given the variation in all these settings for the conditioning sessions (tone-shock, tone-loom and tone alone) it was difficult to standardize the freezing-settings across conditions during the conditioning phase. Hence manual scoring using the open-source program Python Video Annotator (https://pythonvideoannotator.readthedocs.io) was done instead for the conditioning sessions of both experiments, while FreezeScan was used to score freezing during the test sessions aimed at testing learned freezing (first experiment).

Videos obtained from the training and test sessions of the escape experiment were used to analyze various behaviors (particularly displacement and pellet retrieval). Time of pellet retrieval, freezing, scanning, and the time spent in the shelter were scored manually. The position in the arena of each animal for the duration of training and test was determined using Bonsai visual programming language (Lopes et al., 2015). Information from manual scoring and automated scoring was combined with custom Python (Spyder v3.6) scripts and further analyzed.

### Statistical analysis

Statistical analyses were performed using the PRISM 8 software (Graphpad) and Python. For the conditioned freezing experiment, differences between groups were investigated using a *Mann-Whitney U* test. Within subject changes in freezing from baseline to stimulation period were investigated with a *Wilcoxon signed-rank* test. In the escape experiment, a *Kruskal-Wallis* analysis with post hoc Dunn’s test for multiple comparisons was done to investigate the difference between time spent freezing during conditioning of the three groups. Within-subject changes in time spent freezing were again analyzed with a *Wilcoxon signed-rank* test. Statistical significance was accepted at p-value<0.05 for all tests. Regarding the pellet retrieval in the escape experiment, comparisons between the tone-shock, tone-loom and tone-alone groups were done using a *Kaplan-Meier* test for survival (of the pellet in this case).

## Supporting information

video 1 - shock

video 2-loom

video 3-loom

video 4-tone-shock

video 5-tone-loom

video 6-tone

## Acknowledgements

This work was developed with the support from the Rodent platform and the Scientific Software platform at Champalimaud, and the research infrastructure Congento LISBOA-01-0145-FEDER-022170. We want to thank João Frazão for his support with Bonsai, and Gil Costa for the illustrations in the figures. We thank the Moita lab members and Andreia Cruz for helpful discussions and comments on the manuscript.

## Data availability Statement

All behavioral videos are stored on hard drives at the Champalimaud Centre for the Unknown and will be shared freely upon request. The tracked data, python code, bonsai code, and all data used to create the graphs will be shared on an online platform.

## Author contributions

MH and MAM designed the research, discussed the results and wrote the paper. MH performed all experiments and data analysis.

