## supplementary figures for "Looming stimuli reliably drive innate, but not learned, defensive responses in rats"

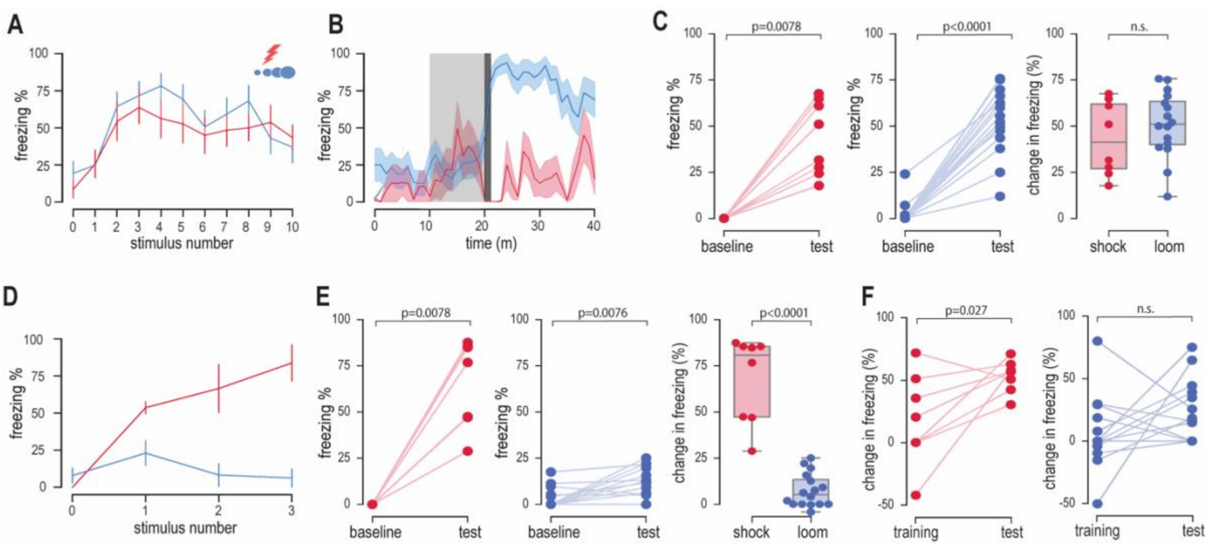


**Supplemental figure 1. Percent time freezing during ten-second tones.** Throughout the figure red represents tone-shock condition (n=8) and blue the tone-loom condition (n=16). **(A)** Freezing during tone presentations of the conditioning session. Line graph showing mean freezing for each of the ten tones (mean +/- S.E.M.). The x-axis shows the number of tones, with 0 being the ten seconds immediately preceding the first tone (see seconds 0-10 in B for the trace during this time period). **(B)** Freezing during 40s around the first loom presentation. Line graph showing freezing averaged across rats (shaded red and blue areas represent the S.E.M.). Each data point corresponds to freezing averaged over 1s epochs. Light grey shade indicates the time of tone presentation, while dark grey indicates the presentation of either shock or loom. **(C)** Left and middle panel: percent time freezing per animal (for tone-shock and tone-loom conditions respectively) during the baseline and the tone presentations (average of all ten tone presentations). To approximate the estimate of freezing levels during baseline to that of freezing during tone presentations, we measured freezing during ten 10s time-windows of the baseline and then averaged across the ten time-windows (each 10s time-window was taken at the end of each minute of the baseline, which lasted 10m in total). Right panel: increase in freezing percentage for each individual in the tone-shock (red) and tone-loom condition (blue) respectively (average freezing during tone - average freezing during baseline). **(D)** and **(E)** same as in **(A)** and **(C)** for freezing during tone presentations of the recall test session. **(F)** Change in freezing to the first tone, during both conditioning and test, is shown for each animal individually. The change in freezing is calculated by subtracting freezing during the ten seconds directly preceding the tone (see point 0 on x-axis in **(A)** and **(D)**) from freezing during the tone (point 1 on x-axis in **(A)** and **(D)**).

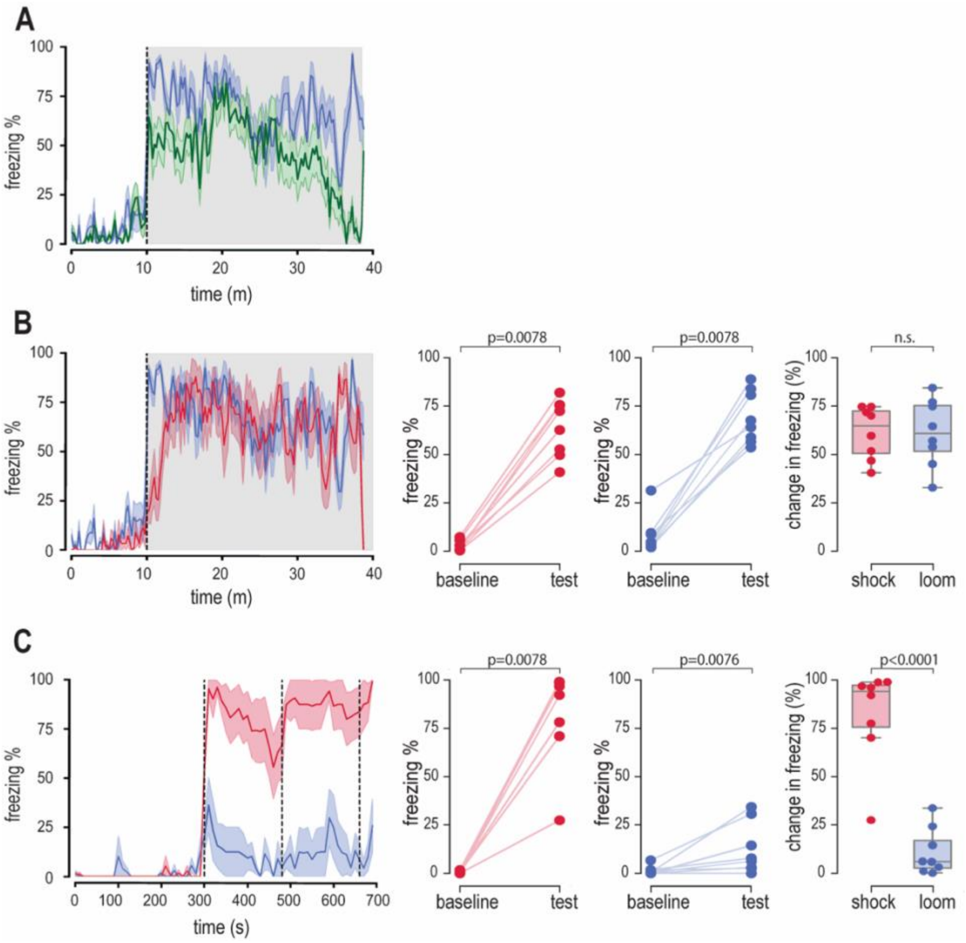


**Supplemental figure 2. The rats in tone-loom condition that freeze to a high extend throughout the conditioning still freeze significantly less during recollection test than rats in tone-shock condition.**

**(A)** Line graph showing freezing per ten-second epoch (average across rats +/- S.E.M.) throughout the conditioning session (grey shade indicates period of tone-loom presentations). Blue and green correspond to high (n=8) and low freezers (n=8) in the tone-loom group, respectively. Rats were categorised as high freezers if they froze >50% of the last minute of the training session. **(B)** Left panel: line graph showing freezing per ten-second epoch (average across rats +/- S.E.M.) throughout the conditioning session (grey shade indicates period of tone-loom presentations). Blue and red correspond to tone-loom high freezers (n=8) and tone-shock group (n=8), respectively. The two middle panels: percent time freezing per animal (for tone-shock and tone-loom high freezers, respectively) during the baseline and the period of stimulus presentations (average of the whole 10m baseline period and over the 10m stimulation period). Right panel: change in freezing percentage for each individual in the tone-shock (red) and tone-loom condition high freezer (blue) respectively (average freezing during tone - average freezing during baseline). **(C)** same as for **(B)** for freezing during the recall test session.

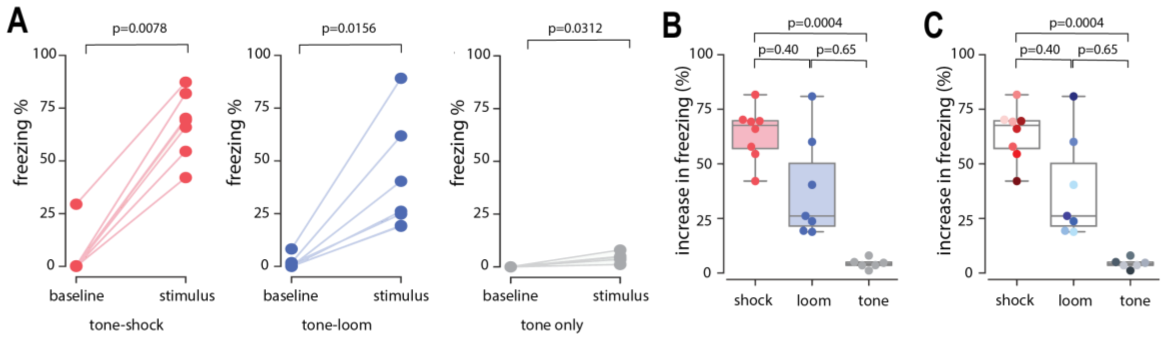


**Supplemental figure 3. Individual freezing increase during conditioning for experiment 2.**

**(A)** Freezing of individuals during exposure to tone-shock (red, n=8), tone-loom (blue, n=7), or tone presentations (gray, n=6). While the amount of increase is very low in the tone group, there is a significant increase in freezing in all three conditions (tone-shock: median increase in freezing: 67.62%, W=36, p=0.0078; tone-loom: median increase in freezing: 26.13%, W=28, p=0.0156; tone: median increase in freezing: 4.09%, W=21, p=0.0312). **(B)** Change in freezing per individual for tone-shock (red), tone-loom (blue), and tone (gray) conditions. Freezing increase upon tone exposure is significantly lower than for the tone-shock condition, while tone-loom exposure resulted in an intermediate freezing increase. (Kruskal-Wallis: p<0.0001 (Kruskal-Wallis statistic=14.53). Dunn's multiple comparisons test: shock vs loom: p=0.3998; shock vs tone: p=0.0004; loom vs tone: p= 0.649). **(C)** Same as in **(B)** with individual data points color-coded by pellet retrieval time (lighter shades for faster retrieval), showing no clear relationship between freezing level and time to retrieve the pellet, regardless of condition.

**Videos – examples of rat behavior during training, conditioning and test.**

**(Video 1) Example of freezing during tone-shock conditioning and tone test after conditioning.** First segment shows an example of the innate response to the first tone-shock pairing of the training session. Second segment shows the learned freezing response of the same rat triggered by shock-associated tone.

**(Video 2) Example of freezing during tone-loom conditioning and absence of freezing in response to the tone test after conditioning.** First segment is an example of the innate defensive freezing response to the first tone-loom pairing of the training session. Second segment shows an example animal's lack of freezing to a loom-associated tone, while displaying risk assessment behaviors.

**(Video 3) Example of pellet retrieval during training and escape in response to the looming stimulus.** First segment shows a rat retrieving the third pellet (furthest from the shelter) of the second pellet-retrieval training session. Second segment shows an escape response of that rat to a looming stimulus that was presented when the rat reached the middle of the runway (before it reached the pellet).

**(Video 4) Example of pellet retrieval during training, freezing during conditioning and escape in response to the shock-associated tone.** First segment shows a rat retrieving the third pellet (furthest from the shelter) of the second pellet retrieval training session. Second segment shows the same rat freezing in response to the last tone-shock pairing during conditioning. Final segment shows the escape response to presentation of the shock-associated tone in the runway.

**(Video 5) Example of an escape responses to loom-associated tone and of a successful pellet retrieval despite the presentation of the loom-associated tone.** Example 1) First segment shows a rat retrieving the third pellet (furthest from the shelter) of the second pellet retrieval training session. Second segment shows the same rat freezing in response to the last tone-loom pairing during conditioning. Third segment shows this rat turning back to the shelter, before pellet retrieval, in response to the loom-associated tone in the runway. Example 2) Fourth and fifth segments show a different animal retrieving the third pellet during the second pellet-retrieval training session and the freezing response during conditioning, respectively. Final segment shows that same rat retrieving the pellet after the loom-associated tone was played.

**(Video 6) Example of an escape responses to a neutral tone and of a successful pellet retrieval despite the presentation of the neutral tone.** Segments 1-6 as described above, except the tone is neutral instead of loom-associated.
